# ClearScope: a fully integrated light sheet theta microscope for sub-cellular resolution imaging without lateral size constraints

**DOI:** 10.1101/2024.08.15.608141

**Authors:** Matthew G. Fay, Peter J. Lang, David S. Denu, Nathan J. O’Connor, Benjamin Haydock, Jeffrey Blaisdell, Nicolas Roussel, Alissa Wilson, Sage M. Aronson, Paul J. Angstman, Cheng Gong, Tanvi Butola, Orrin Devinsky, Jayeeta Basu, Raju Tomer, Jacob R. Glaser

## Abstract

Three-dimensional (3D) ex vivo imaging of cleared intact brains of animal models and large human and non-human primate postmortem brain specimens is important for understanding the physiological neural network connectivity patterns and the pathological alterations underlying neuropsychiatric and neurological disorders. Light-sheet microscopy has emerged as a highly effective imaging modality for rapid high-resolution imaging of large cleared samples. However, the orthogonal arrangements of illumination and detection optics in light sheet microscopy limits the size of specimen that can be imaged. Recently developed light sheet theta microscopy (LSTM) technology addressed this by utilizing a unique arrangement of two illumination light paths oblique to the detection light path, while allowing perpendicular arrangement of the detection light path relative to the specimen surface. Here, we report development of a next-generation, fully integrated, and user-friendly LSTM system for rapid sub-cellular resolution imaging uniformly throughout a large specimen without constraining the lateral (XY) size. In addition, we provide a seamlessly integrated workflow for image acquisition, data storage, pre- and post-processing, enhancement, and quantitative analysis. We demonstrate the system performance by high-resolution 3D imaging of intact mouse brains and human brain samples, and complete data analysis including digital neuron tracing, vessel reconstruction and design-based stereological analysis in 3D. This technically enhanced and user-friendly LSTM implementation will enable rapid quantitative mapping of molecular and cellular features of interests in diverse types of very large samples.

## INTRODUCTION

Three-dimensional (3D) ex vivo, whole-brain imaging using transgenic and non-transgenic animal models, as well as large human and non-human primate postmortem brain specimens, holds immense potential to gain novel insights into both physiological neural network connectivity patterns and pathological alterations of connectivity associated with neuropsychiatric and neurological conditions.^1-3^

Light-sheet microscopy (LSM) has proven to be the most effective imaging modality for whole-brain datasets compared to confocal and multiphoton microscopy because it provides a high resolving power in 3D while enabling more rapid data collection and lower phototoxicity.^4^ LSM yields the best results when combined with tissue clearing methods (including CLARITY^5-7^, uDISCO^8^, SeeDB^9^, Sca*l*e^10^ and Binaree Tissue Clearing^11^), which make it possible to optically access intact tissue while preserving the brain’s molecular and structural architecture, in particular all the neuronal connections.^3^

While more than 30 LSM technologies are reported in the literature^4^ most are unavailable, commercially or otherwise.

The recently developed light sheet theta microscopy (LSTM) technology^12^ uses a unique arrangement of two illumination light paths oblique to the detection light path, with perpendicular arrangement of the detection light path relative to the specimen. This unique arrangement serves as the basis of the following capabilities of the LSTM technology that are not all available with any of the LSM systems that are currently available commercially or otherwise: (i) imaging thick specimens of unlimited lateral (XY) size up to a maximum depth (Z) range that is constrained only by the working distance of the detection objective used; (ii) illuminating the specimen from two different angles simultaneously, thereby always providing excitation light that reaches regions behind opaque structures in the specimen; (iii) use of different objectives in the illumination light paths and the detection light path (with different magnification and different numerical aperture) for optimal imaging performance, and (iv) the possibility to use objectives with different refractive indices (depending on the tissue clearing method used) and from different providers in order to enable users to make use of innovations in LSM objective technology that may be developed and marketed in the future.

Dr. Tomer and his team successfully established proof of concept and demonstrated that the development of the LSTM technology represents clear progress beyond the state-of-the-art.^12^ This project aimed to further develop the LSTM technology, including creating and testing (i) a new, optimized design, (ii) a novel chamber that contains the investigated specimen and the immersion medium as well as (iii) a novel detection objective changer, (iv) novel image acquisition software including features for adaptive refractive index mismatch correction and (v) novel software for stitching image stacks acquired with the LSTM technology to create composite 3D images without the need to downsample the image information.

The ultimate aim of this project was (i) to develop a novel, fully integrated light sheet theta microscope (ClearScope^®^) that enables sub-cellular resolution imaging of specimens with unconstrained lateral (XY) size with a depth (Z) range that is constrained only by the working distance of the detection objective used, and (ii) to provide a seamless workflow, integrating image acquisition, data storage, image post-processing and enhancement, and quantitative analysis in a way that performing light sheet microscopy can be done without any expertise in software programming and/or microscope hardware assembly

## RESULTS

### Microscope hardware design

The ClearScope and its predecessor, the original LSTM system^12^, are shown in Figure 1 (the abbreviations provided in the following text refer to Figs 1C and 2A-D). Laser (l) light passes through a laser collimator (lc), first iris (i_1_), electrically tuneable lens (etl), adjustable slit (as), cylindrical lens (cl), galvo scanner (gs), f-theta scan lens (sl), second iris (i_2_), tube lens (tl) and illumination objective (io). During imaging with two light sheets (ls_1_, ls_2_), the specimen chamber (sc) containing the specimen (s) is filled with immersion oil with appropriate refractive index. In the detection light path a variety of objectives with long working distance (WD) can be used, including a MBF Bioscience modified UPLFLN4XPH objective (4x/0.13 NA; WD = 17 mm; Olympus, Tokyo, Japan), XLPLN10XSVMP objective (10x/0.6 NA ∞, WD = 8 mm; Olympus) and CFI90 20XC Glyc objective (20x/1.0 NA ∞, WD = 8.2 mm; Nikon, Tokyo, Japan). Note that in the ClearScope the WD of the XLPLN10XSVMP objective is reduced from 17 mm to 12 mm due to the use of a correction lens. The latter is required to use the UPLFLN4XPH air objective in the presence of immersion oil (as implemented in the ClearScope).

**Fig. 1.**
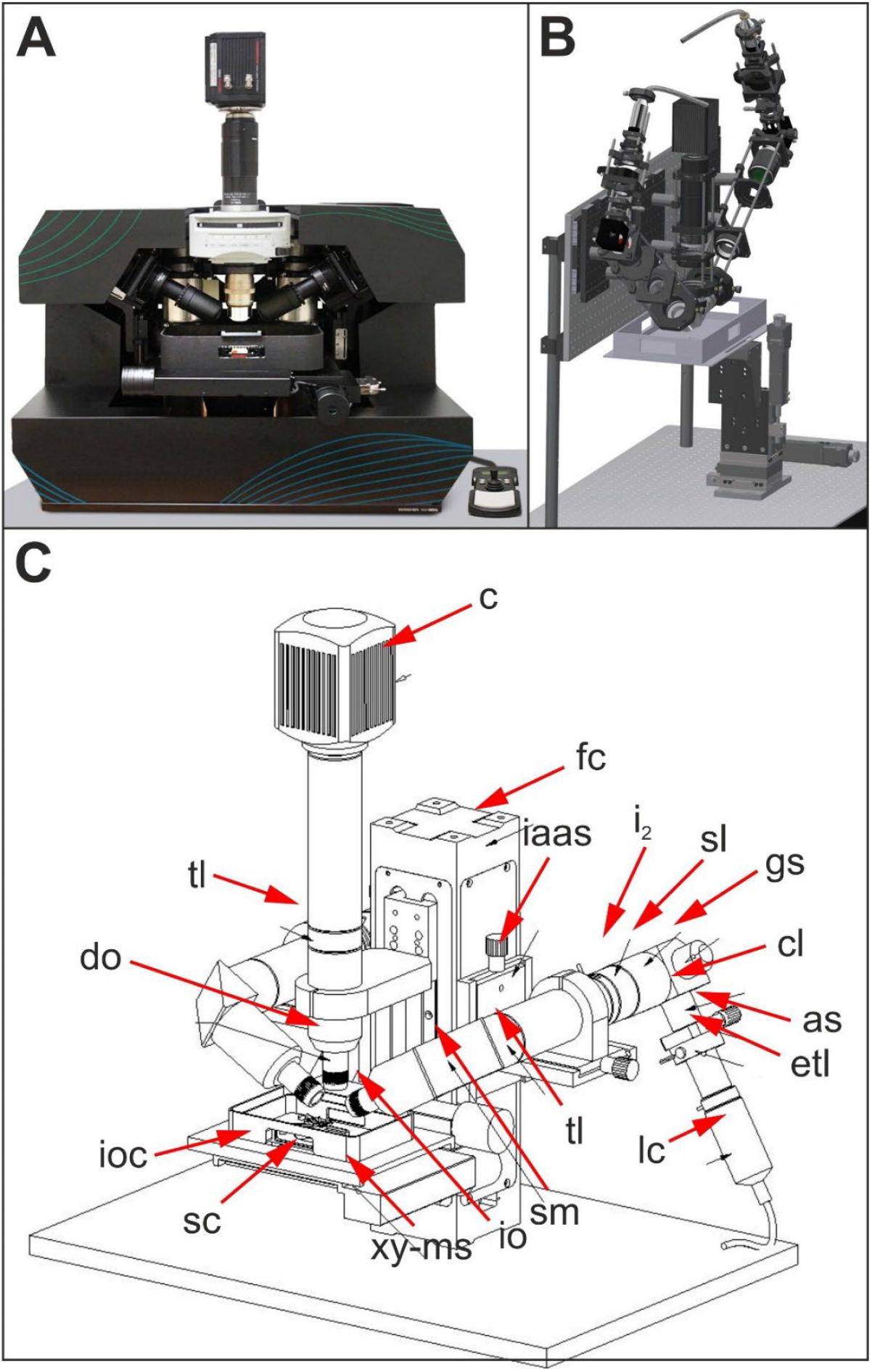
(**A**) Production version of the ClearScope developed in this project. (**B**) 3D CAD sketch of the original light sheet theta microscopy system.^12^ (**C**) 3D CAD drawing of the ClearScope. Abbreviations: as, adjustable slit; c, camera; cl, cylindrical lens; do, detection objective; etl, electrically tuneable lens; fc, focus column; gs, galvo scanner; i2, second iris; iaas, illumination light arm alignment slide; io, illumination objective; ioc, immersion oil chamber; lc, laser collimator; sc, specimen chamber; sl, f-theta scan lens; sm, steering mirror; tl, tube lens; xy-ms, XY motorized stage. Details are in the text. Panel B was taken without modification from^12^ published under the CC BY 4.0 license (https://creativecommons.org/licenses/by/4.0/).

**Fig. 2.**
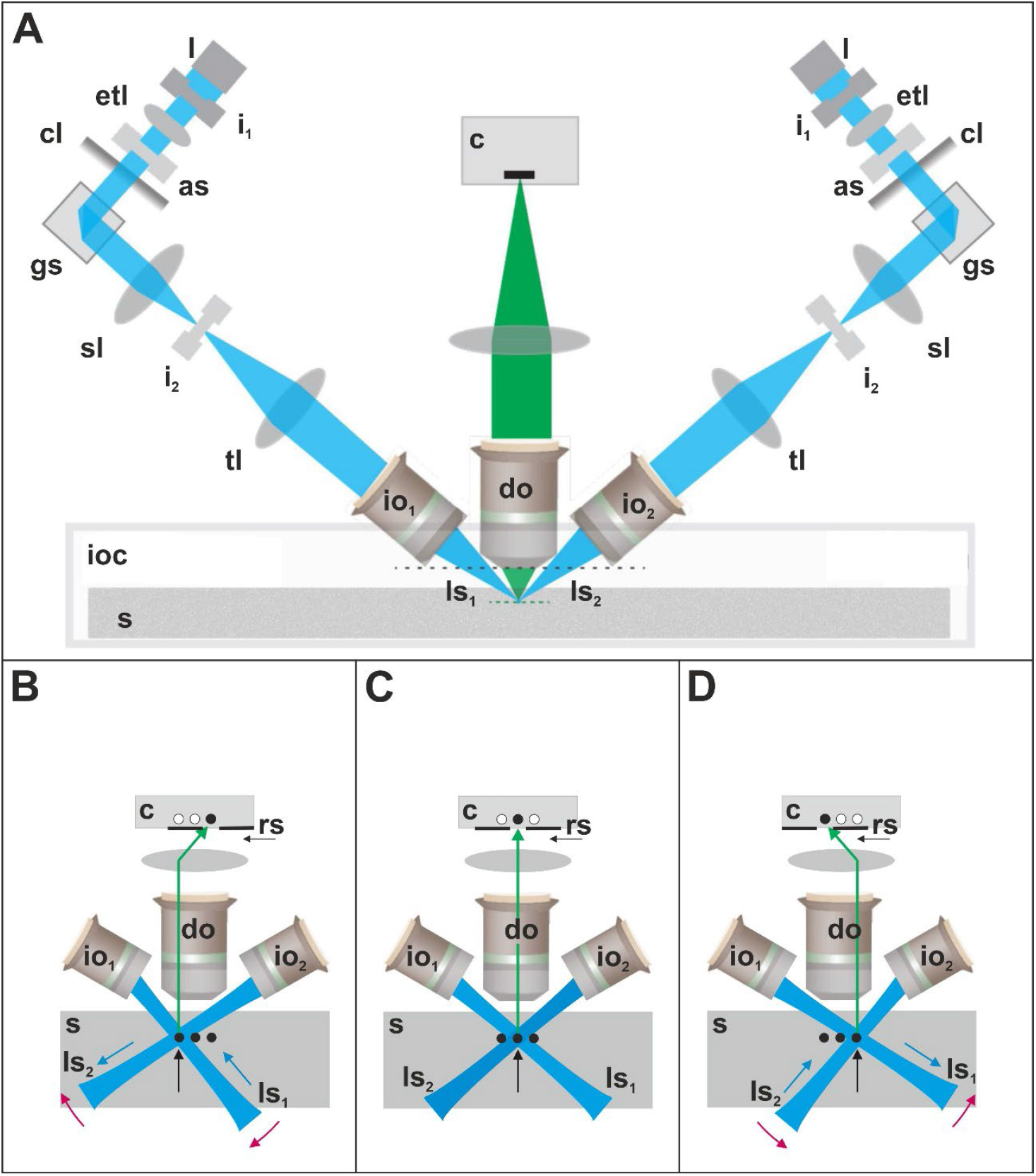
Principle of imaging with the ClearScope developed in this project. Abbreviations: as, adjustable slit; c, camera; cl, cylindrical lens; do, detection objective; etl, electrically tuneable lens; gs, galvo scanner; i_1_, first iris; i_2_, second iris; io_1_, first illumination objective; io_2_, second illumination objective; ioc, immersion oil chamber; l, laser; ls_1_, first light sheet; ls_2_, second light sheet; rs, rolling shutter; s, specimen; sl, f-theta scan lens; tl, tube lens. Details are in the text. Panel A was modified from^12^ published under the CC BY 4.0 license (https://creativecommons.org/licenses/by/4.0/).

The principle of imaging with the ClearScope is shown in Figure 2, showing a single field-of-view (FOV) at different positions of two light sheets (ls_1_, ls_2_) in order to sequentially image three lines within a certain focal plane inside the specimen (s) (indicated by black dots and black arrows in Fig. 2B-D) using a rolling shutter (rs). Compared to imaging the line in the middle of the FOV (Fig. 2C), imaging the line on the left requires turning both light sheets clockwise (red arrows in Fig. 2B) and move ls_1_ “into” the first illumination objective (io_1_) as well as ls_2_ “out” of the second illumination objective (io_2_) (blue arrows in Fig. 2B). Conversely, imaging the line on the right requires both light sheets to turn counterclockwise (red arrows in Fig. 2D) and move ls_1_ “out” of io_1_ as well as ls_2_ “into” io_2_ (blue arrows in Fig. 2D). Turning the light sheets clockwise and counterclockwise is achieved by the galvo scanners; moving the light sheets into and out of the illumination objectives is achieved by the electrically tuneable lenses.

Figure 3 shows the specimen holding assembly of the ClearScope. A rectangular aluminium tub (100 x 200 x 44 mm (W x L x H)) mounted on top of a stage insert serves as immersion oil chamber (Fig. 3A; the asterisk indicates the immersion oil). Within the center of the tub is the specimen chamber (asterisk in Fig. 3A) which consists of a magnetic coupler fixed to the bottom of the tub (not shown in Fig. 3A), a chamber holder (Fig. 3B) and a removable chamber that can accommodate a whole mouse or rat brain (arrow in Fig. 3B).

**Fig. 3.**
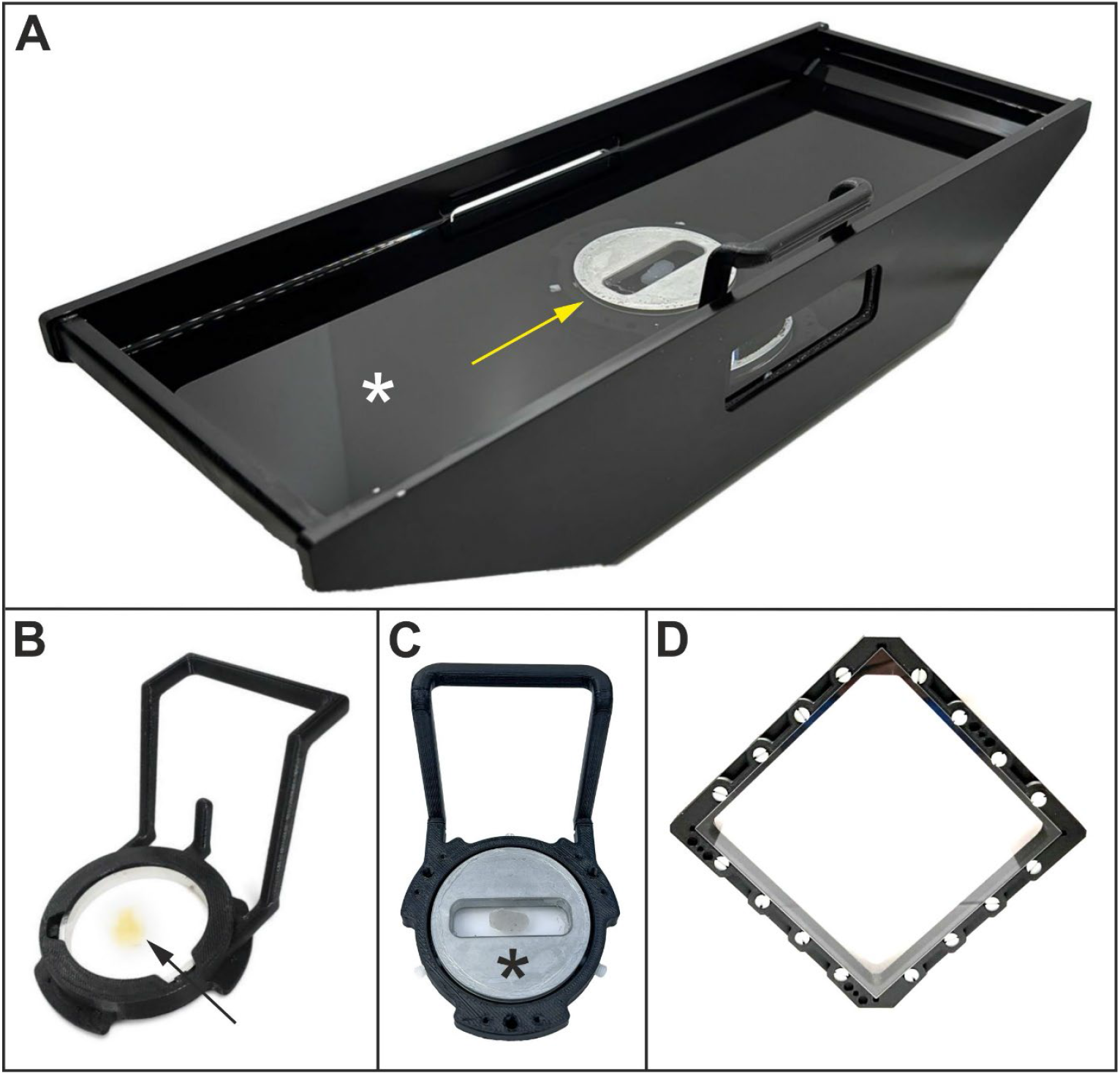
The specimen holding assembly of the ClearScope developed in this project. Details are in the text.

The chamber itself consists of two round coverslips (thickness no. 1; 50 mm diameter), separated by silicon gaskets which provide an adjustable height of the chamber to accommodate rodent brains of different sizes. All of this are contained within an aluminium ring with a threaded internal collar which creates a seal of the coverslips against the silicon gaskets. For imaging smaller specimens, a low-volume specimen chamber (asterisk in Fig. 3C; also shown in Fig. 3A) can be inserted into the chamber holder.

The XL specimen chamber is shown in Figure 3D. The interior dimensions of the XL specimen chamber are 144 x 144 x 8 mm (W x L x H); it is a symmetrical design that can be used with either side facing up, enabling imaging of one side followed by imaging of the other side if desired (for use of the XL specimen chamber a XL immersion oil chamber was developed that is not shown in Fig. 3). The frames have a channel to hold vacuum grease that seals them to the specimen windows, and the XL specimen chamber assembly is secured to the immersion oil chamber using the same magnetic/mechanical detent holder mechanism used for other ClearScope specimen holders (c.f. Fig. 3A).

Figure 4 shows the detection objective changer (DOC) of the ClearScope and its working principle. The DOC enables to flexibly change the detection objective even during examination of a specimen according to the users’ imaging needs. Repeatability tests demonstrated that the position variability is <3 µm, which can effectively be corrected with the ClearScope system software.

**Fig. 4.**
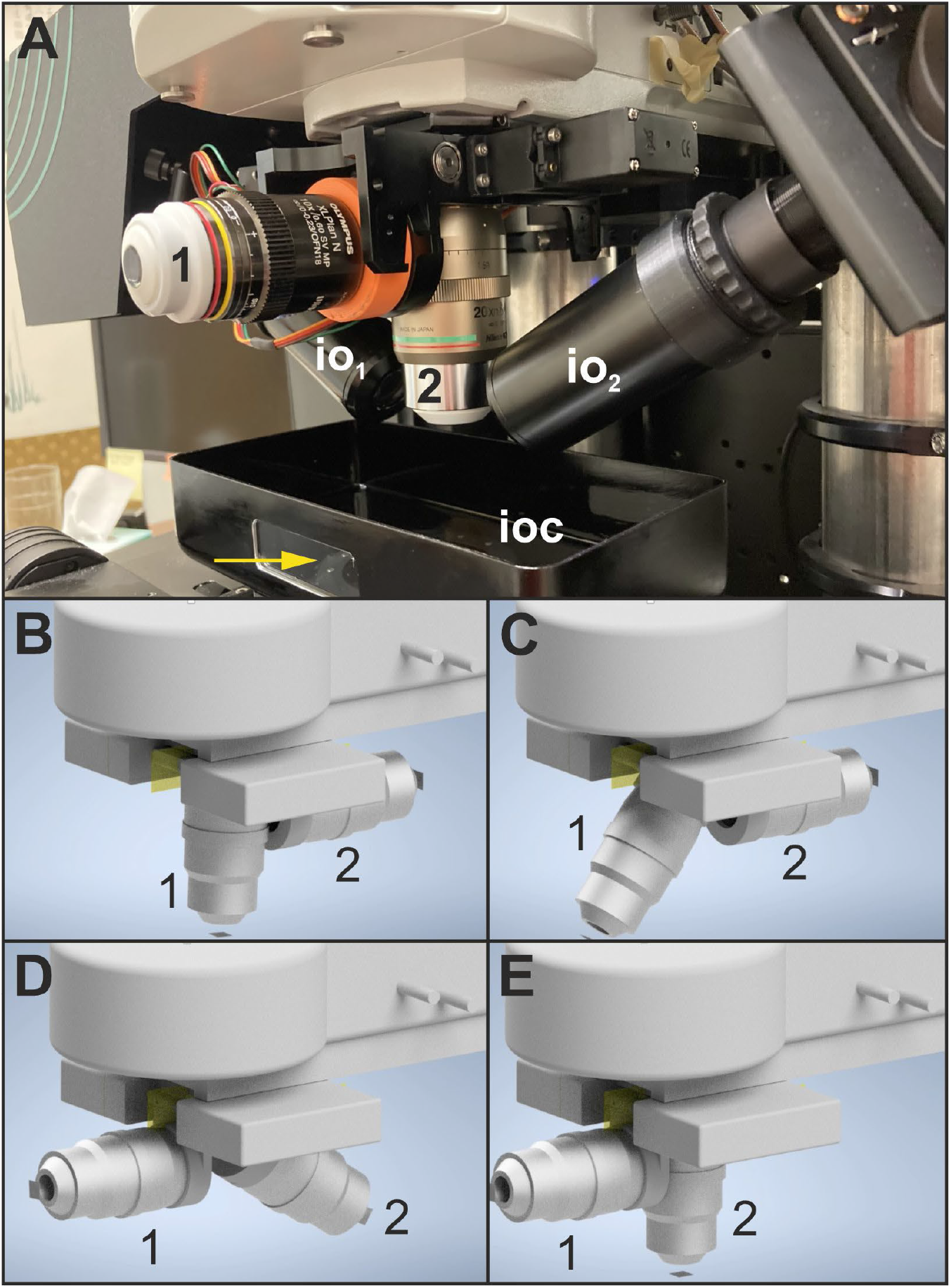
(**A**) The detection objective changer (DOC) of the ClearScope developed in this project. Abbreviations: 1, first detection objective (here: XLPLN10XSVMP oil; NA = 0.6; WD = 8 mm; Evident Corp., Tokyo, Japan); 2, second detection objective (here: CFI90 20XC Glyc; NA = 1.00; Nikon); io_1_, first illumination objective; io_2_, second illumination objective; ioc, immersion oil chamber. The arrow points to a window in the immersion oil chamber through which the light sheets can be observed (c.f. Fig. 5). (**B**-**E**) Working principle of the DOC. (**B**) System is ready for lens change; stage is in load position. (**C**) The first detection objective (1 in (A-E)) begins to swing away from the active position. (**D**) Swing continues. (**E**) Swing complete; the second detection objective (2 in (A-E)) is poised in position.

### Microscope software design

Image acquisition software was created to control all components of the ClearScope, synchronizing the camera exposure window, galvo scanning and ETL focus to maintain the thinnest (focused) part of the light sheets, which enables optimal conditions for image acquisition (Fig. 5).

**Fig. 5.**
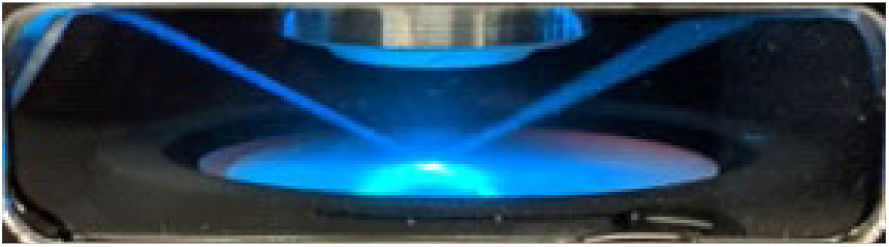
Visualization of the two light sheets of the ClearScope developed in this project, viewed through the window in the immersion oil chamber (arrow in Fig. 4A). Imaging is performed along an “illumination line” at the position where the thinnest parts of both light sheets overlap (minimum thickness 3.2 µm).

Synchronization of the galvo scanning and ETL focus adjustments for 2-axis scanning was optimized using polynomial regression analysis of 7 calibration points across the FOV. This method accounts for nonlinearities in ETL focus behavior, improves overall image quality, and enables utilization of larger FOV objectives and cameras. *Intelligent Refractive Index Compensation* (IRIC) was developed to enable use of the ClearScope with specimens cleared using a wide array of techniques, further increasing system flexibility. For each color channel, IRIC automatically adjusts the position and focus of the light sheets as a function of tissue depth, enabling a single detection objective to image tissues cleared with several different protocols, including CLARITY^5-7^, uDISCO^8^, SeeDB^9^, Sca*l*e^10^ and Binaree Tissue Clearing^11^.

Although the original IRIC implementation works well for smaller samples, (e.g., mouse brain, organs, etc.), it turned out that a single correction through Z applied to larger samples (e.g., human brain slabs) was insufficient. To remedy this, a 3D mapping version of the IRIC was implemented that uses correction data from multiple XY locations to compensate for heterogeneity throughout large tissue samples. The user interface is designed to make it easy for users to select appropriate refractive-index configuration settings.

To avoid problems during acquisition of large specimens, a disk-space verification function was developed that estimates the disk space that will be needed for the acquisition and verifies that it is available. This is an important feature because image acquisition often proceeds when the microscope operator is not present.

New post-processing image filters were added to the ClearScope software, including a new flat-field correction and blending algorithm to seamlessly stitch multiple FOVs with uniform illumination appearance, a rolling-ball filter to mitigate the uneven illuminated background, and an unsharpen mask filter to further enhance overall image quality for publications or presentations. Uneven illumination is a phenomenon typically caused by aberrations in the illumination optics. It can introduce considerable bias in image acquisition. In the ClearScope system, we found that illumination variations can be reliably modelled by a linear intensity gain function combined with an additive term. Estimating the distortion model enabled distortions of the illumination to be reversed to generate corrected images. The ClearScope software utilizes a very flexible, retrospective model estimation method whereas the distortion field is estimated after acquisition using regularized energy minimization^13^. The overall quality of the final image montage is further improved with linear blending to smooth pixel transition in the overlapping areas.

The fastest exposure time that was achieved was 45.9 ms per plane. In light sheet mode, the camera employs a rolling shutter that maximizes signal-to-noise ratio, but at a reduced overall image-acquisition rate compared to using the camera in widefield mode. Minimizing image-acquisition time was achieved by overlapping software functions (e.g., setting galvo and ETL information) to coincide with Z movements, decreasing camera-exposure times using the more sensitive Prime BSI Express camera (Teledyne Photometrics, Tucson, AZ, USA) instead of the Orca Fusion Digital CMOS camera (Hamamatsu, Hamamatsu City, Japan) that was originally used, and improving ETL focus during scanning.

An intuitive user-interface was created for both configuration and operation of the DOC and the following 7-channel, single-mode fiber laser systems: Cobolt C-Flex C8 (Hübner Photonics Inc., San Jose, CA, USA) and iChrome FLE (TOPTICA Photonics Inc., Pittsford, NY, USA). Experiments with several collimation optical designs demonstrated that single-mode fiber laser systems produced thinner light sheets and higher axial resolution than multi-mode fiber laser systems (Lumencor Celesta and Ziva (Lumencor, Beaverton, OR, USA); and the LDI (89 North, Williston, VT, USA)).

The ClearScope automatic tissue-detection functionality is used, in conjunction with automatic stage movement and the DOC, to scan and acquire images at relatively low Z-plane resolution. An intuitive *Tissue Scanning Workflow* (TSW) was developed and implemented to easily guide users through the multi-resolution imaging operation. Once the low-resolution scan is complete, a preview of the acquired image is displayed. ClearScope users can then specify regions of interest (ROIs) in the preview image by drawing contours to indicate the areas to be imaged at higher resolution. After switching the detection objectives either manually or using the DOC, the user repeats the TSW to acquire the higher resolution images from the previously identified ROIs. Upon completion of the high-resolution scan, users are presented with a preview of the high-resolution images and may select from several tools, e.g., 3D stitching, flatfield correction, etc., to maximize image quality and utility for their specific research purposes.

The user interface of the ClearScope software for image acquisition and microscope control is shown in Figure 6.

**Fig. 6.**
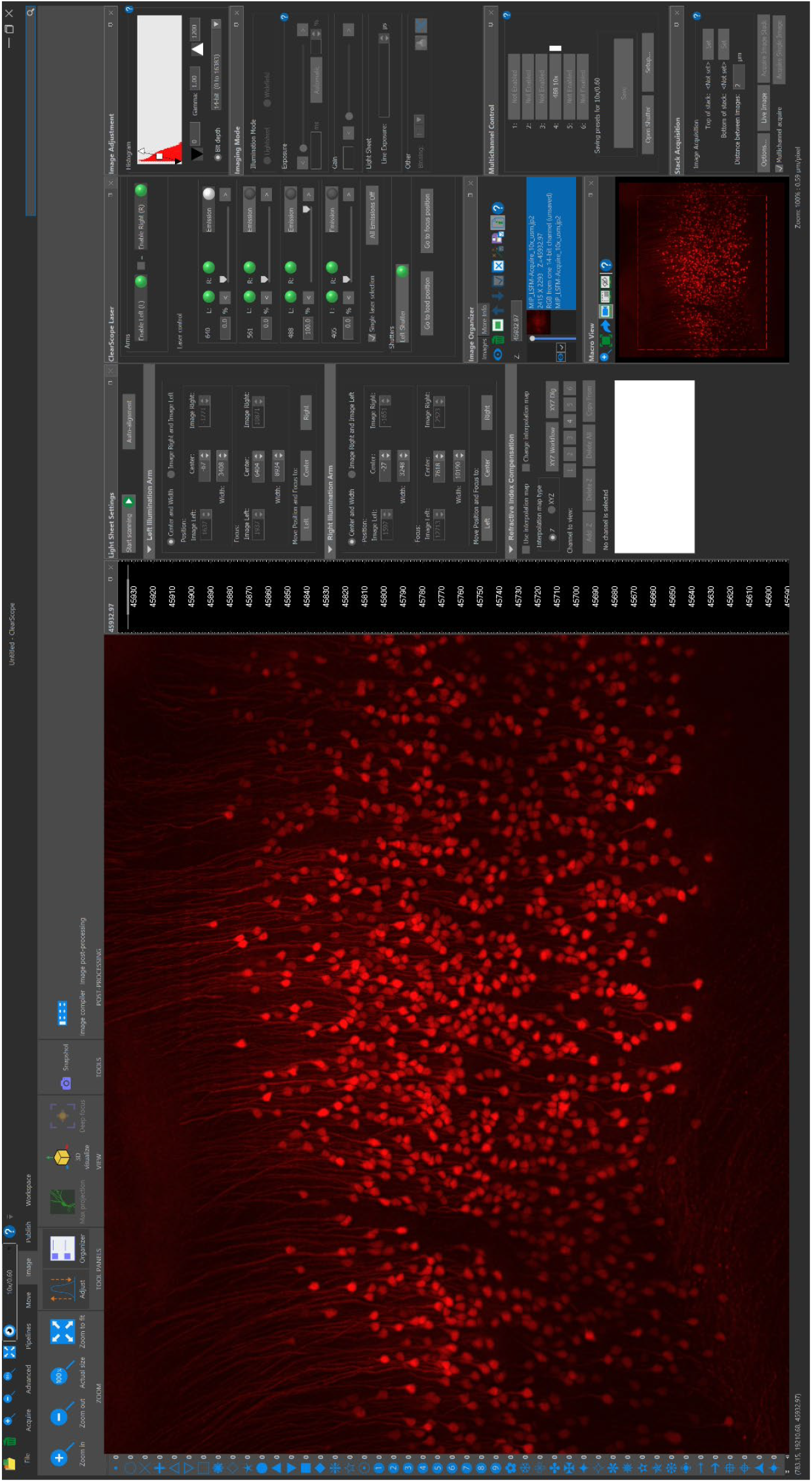
The user interface of the ClearScope software for image acquisition and microscope control.

### Assessment of the image quality

An estimate of the point spread function (PSF) of the ClearScope was generated by using a XLPLN10XSVMP objective (10x/0.6 NA ∞, WD = 8 mm; Olympus) to acquire 36 3D images of the same FOV from a slide containing fluorescent microspheres (diameter, 0.5 µm; Focal Check Fluorescence Microscope Test Slide #5; ThermoFisher Scientific, Waltham, MA, USA) (Fig. 7). Imaging was performed using a 561 nm excitation laser and Prime BSI Express camera (2048 x 2048 pixels; rolling shutter; Teledyne Photometrics). For this optical configuration and a light sheet axial thickness of 4 µm, the axial resolution limit is 4 µm. Images were acquired with an axial sampling of 1.5 µm, satisfying the Nyquist criterion. All 36 FOVs were registered using Big Stitcher^14,15^ and then summed in Fiji^16,17^ to improve image signal-to-noise ratio before making full width at half maximum (FWHM) measurements in Fiji^16,17^ via its intensity profiling tool in the corners and center of the summed FOV.

**Fig. 7.**
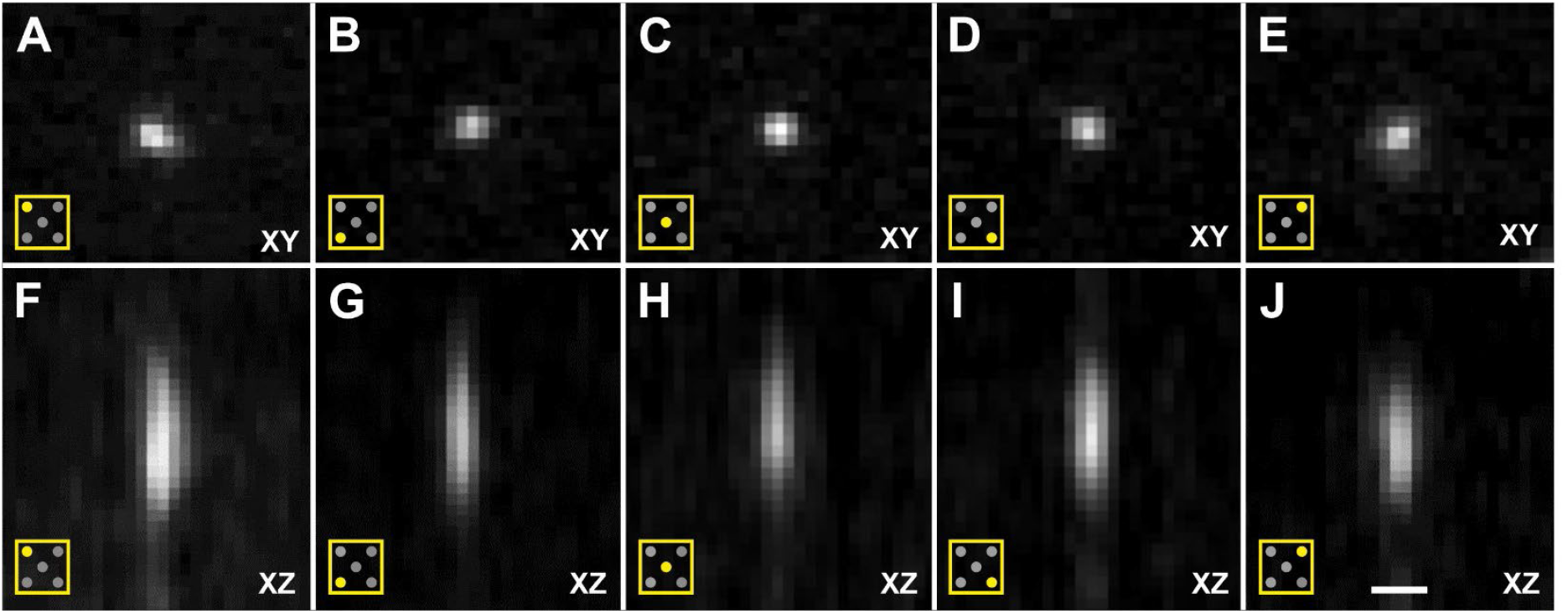
Cross-sections of image stacks generated using Orthogonal Views in FIJI^16,17^ (22 images each; 0.589 µm/pixel in XY; distance between images in Z, 1.5 µm; single image plane shown in XY) of a slide containing fluorescent microspheres (diameter, 0.5 µm; Focal Check Fluorescence Microscope Test Slide #5; ThermoFisher Scientific), captured using a ClearScope (developed in this project) with a XLPLN10XSVMP objective (10x/0.6 NA ∞, WD = 8 mm; Olympus), 561 nm excitation laser and Prime BSI Express camera (Teledyne Photometrics). The panels show XY cross-sections (**A**-**E**) as well as XZ cross-sections (**F**-**J**) of the upper-left quarter of the field of view (FOV) (A,F), the lower-left quarter of the FOV (B,G), the center of the FOV (C,H), the lower-right quarter of the FOV (D,I) and the upper-right quarter of the FOV (E,J) as indicated by the pictograms in the lower left corner in Panels (A-J). These image stacks were used to determine the point spread function of the ClearScope. The scale bar in (J) represents 3 µm in (A-E) and 2,8 µm in (F-J).

Alignment of the image channels was determined on 3D images acquired of the same FOV from a slide containing fluorescent microspheres (diameter, 15 µm; Focal Check Fluorescence Microscope Test Slide #1; ThermoFisher Scientific) (Fig. 8). Imaging was performed using a XLPLN10XSVMP objective (10x/0.6 NA ∞, WD = 8 mm; Olympus), 405nm, 488 nm, 561 nm and 640 nm excitation lasers and Prime BSI Express camera (Teledyne Photometrics). It was found that even when viewed isotropically, the beads appeared spherical, which was due to the very thin light sheets of the ClearScope (minimum thickness 3.2 µm). Furthermore, there was perfect alignment between the different color channels.

**Fig. 8.**
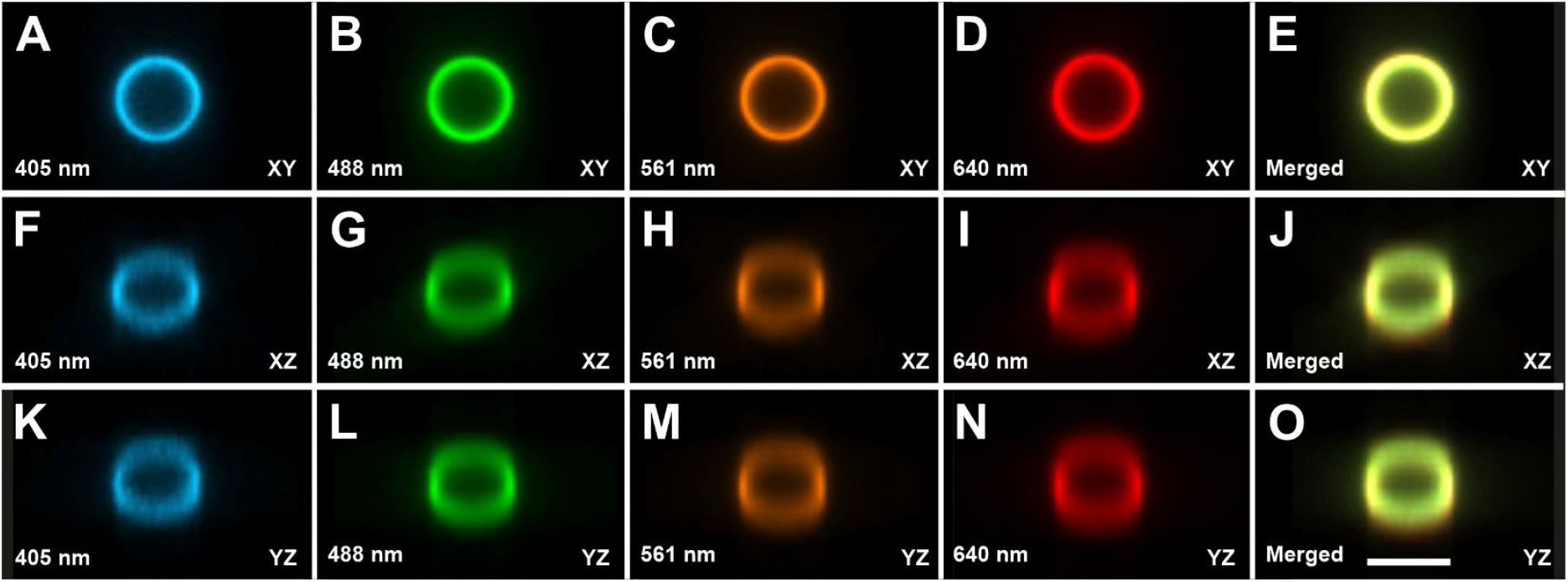
Cross-sections of image stacks generated using Slice Viewer in the 3D environment of the Neurolucida 360 software (MBF Bioscience)^18^ (25 images each; 0.589 µm/pixel in XY; distance between images in Z, 1.5µm; single image plane shown in XY) of a slide containing fluorescent microspheres (diameter, 15 µm; Focal Check Fluorescence Microscope Test Slide #1; ThermoFisher Scientific) emitting light at different wavelengths, captured using a ClearScope (developed in this project) with a XLPLN10XSVMP objective (10x/0.6 NA ∞, WD = 8 mm; Olympus), 405nm, 488 nm, 561 nm and 640 nm excitation lasers (as indicated in the panels) and Prime BSI Express camera (Teledyne Photometrics). The panels show channel-specific (**A**-**N**) XY views (**A**-**D**), XZ views (**F**-**I**) and YZ vies (**K**-**N**) as well as all channels merged (**E**,**J**,**O**). Even when viewed isotropically, the beads appeared spherical, which was due to the very thin light sheets of the ClearScope (minimum thickness 3.2 µm). Furthermore, there is perfect alignment between the different color channels. The scale bar in (O) represents 15 µm.

The axial FWHM was found between 5.45 µm and 6.37 µm, and the lateral FWHM between 1.52 µm and 1.65 µm (Table 1). These data objectively demonstrate that the ClearScope enables imaging at sub-cellular resolution using the 10x objective at all locations within a specimen that is unlimited in lateral (XY) size, and within the 8mm working distance of the objective.

**Table 1.**
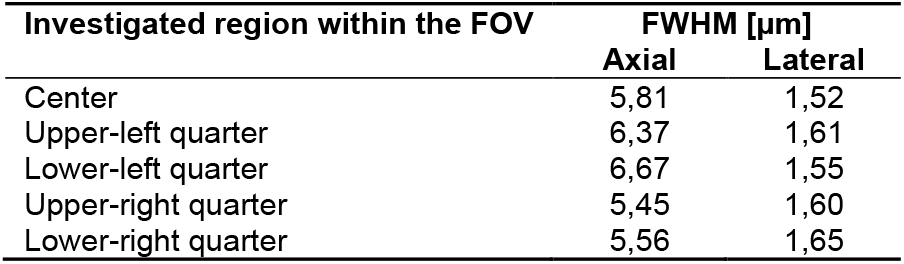
FWHM measurements of the ClearScope developed in this project using a XLPLN10XSVMP objective (10x/0.6 NA ∞, WD = 8 mm; Olympus). Details are in the text.

| Investigated region within the FOV | FWHM [μm] | |
| --- | --- | --- |
|  | Axial | Lateral |
| Center | 5,81 | 1,52 |
| Upper-left quarter | 6,37 | 1,61 |
| Lower-left quarter | 6,67 | 1,55 |
| Upper-right quarter | 5,45 | 1,60 |
| Lower-right quarter | 5,56 | 1,65 |

### Imaging of biological specimens

Various specimens were imaged using a ClearScope; representative examples are shown in Figs 9-13.

**Fig. 9.**
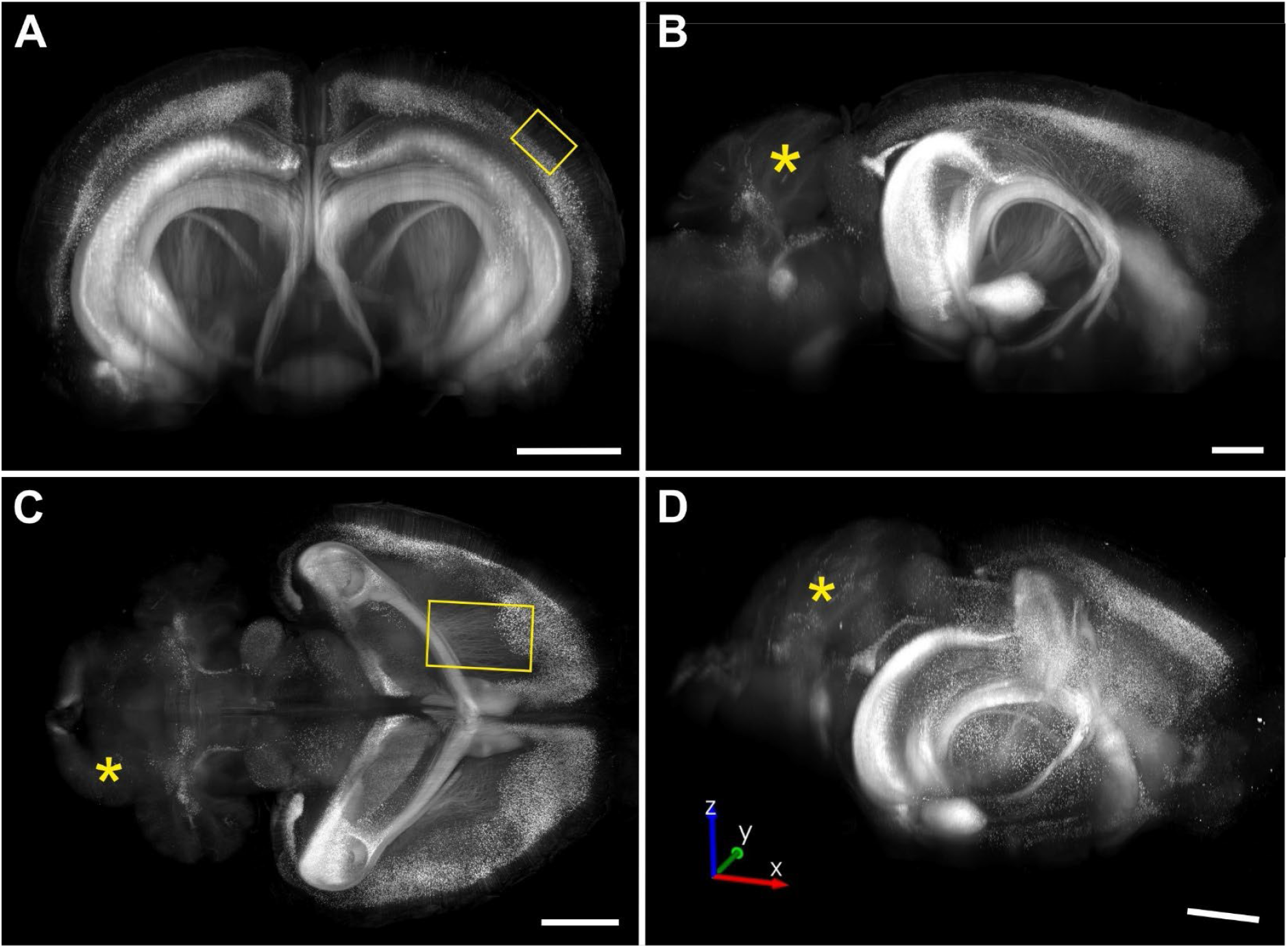
Maximum intensity projections (coronal view in **(A)**, sagittal view in **(B)**, horizontal view in **(C)** and view at an arbitrary angle in (D)) of portions of a 3D image of an entire, intact brain from a transgenic mouse expressing eGFP under control of the Thy-1 promoter acquired using a ClearScope (developed in this project) with a MBF Bioscience modified UPLFLN4XPH objective (4x/0.13 NA; WD = 17 mm; Olympus), 488 nm excitation laser, ZET405/488/561/640mv2 quad band emission filter (Chroma Technologies) and Prime BSI Express camera (Teledyne Photometrics). Note that the weak eGFP signal in the cerebellum (asterisks in (B-D) is expected.19 The brain was provided by Binaree, Inc. (Daegu, Republic of Korea) and cleared with Binaree Tissue Clearing11. The yellow rectangle in (A) indicates the position of the high-magnification detail shown in Fig. 10A, and the yellow rectangle in (C) the position of the high-magnification detail shown in Fig. 11A. The lack of a vertical and horizontal banding pattern demonstrates that this 3D image can be viewed and analyzed from any angle (as shown in (D), which is an indispensable requirement for unbiased, reproducible, digital neuron tracing in 3D. Each scale bar represents 2 mm.

Figure 9 shows maximum intensity projections (MIPs) of portions of a 3D image of an entire, intact brain from a transgenic mouse expressing eGFP under control of the Thy-1 promoter (hereafter, Thy1-eGFP), acquired using a MBF Bioscience modified UPLFLN4XPH objective (4x/0.13 NA; modified WD = 12 mm; Olympus). The lack of a vertical and horizontal banding pattern demonstrates that this 3D image can be viewed and analyzed from any angle (as shown in Fig. 9D), which is an indispensable requirement for unbiased, reproducible, digital neuron tracing in 3D.

Figure 10 shows high-magnification ROI acquisitions of the same mouse brain that is shown in Figure 9, acquired using a XLPLN10XSVMP objective (10x/0.6 NA ∞, WD = 8 mm; Olympus). Figure 10A demonstrates unequivocal identification of cell bodies of individual neurons in the cerebral cortex as well as the apical dendrites of these neurons. Furthermore, Figure 10B shows a cell process that passes tangentially through the cerebral cortex of the same brain and could be traced over a distance of 650 µm in the image shown. In addition, Figure 11A demonstrates cell processes in the subcortical white matter in the 3D image of a mouse brain that is shown in Figure 9C that can be individually traced.

**Fig. 10.**
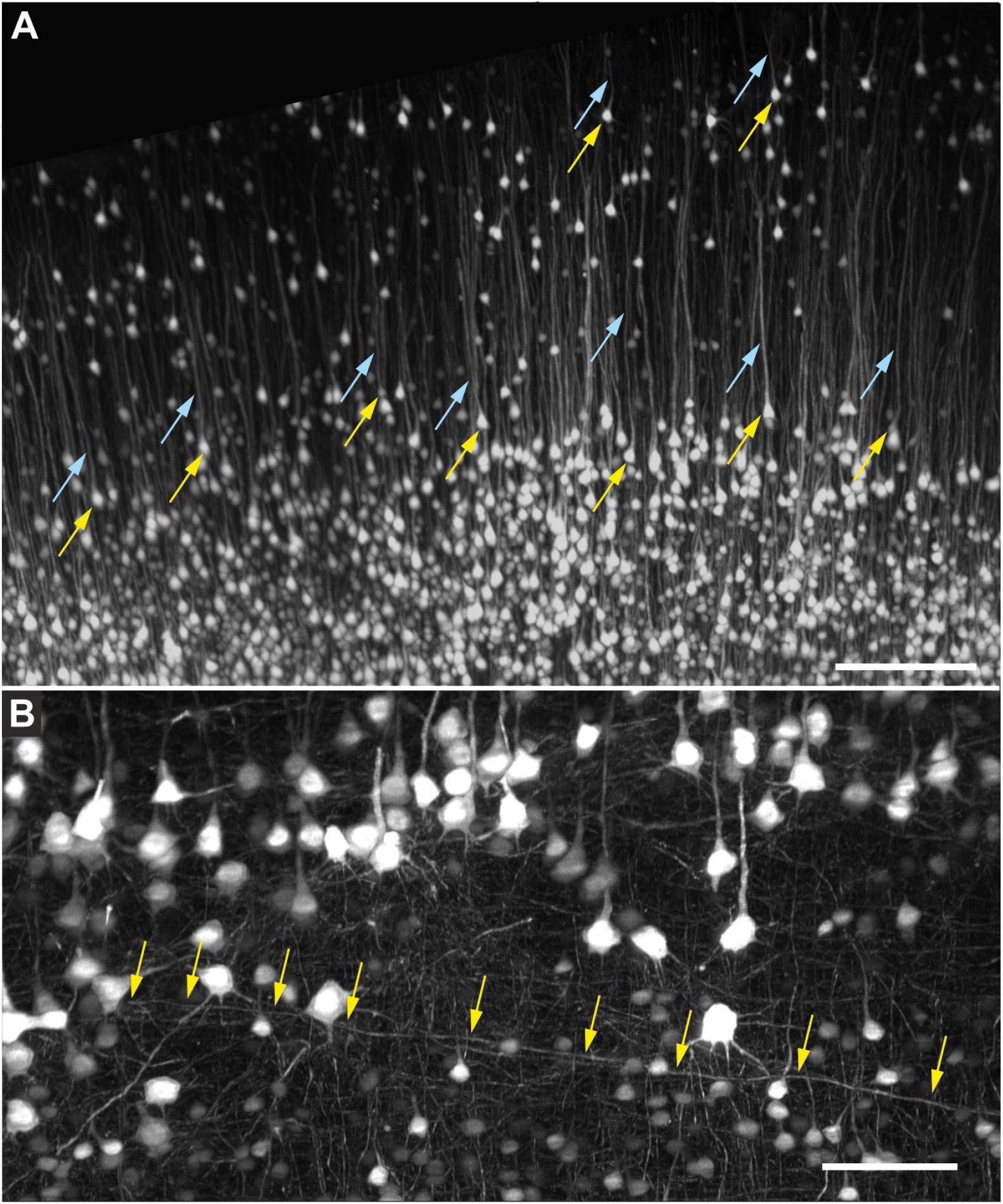
(**A**) High-magnification region of interest acquisition of the region indicated by the yellow rectangle in Fig. 9A of an entire, intact brain from a transgenic mouse expressing eGFP under control of the Thy-1 promoter, acquired using a ClearScope (developed in this project) with a XLPLN10XSVMP objective (10x/0.6 NA ∞, WD = 8 mm; Olympus), 488 nm excitation laser, ZET405/488/561/640mv2 quad band emission filter (Chroma) and Prime BSI Express camera (Teledyne Photometrics). The yellow arrows indicate cell bodies of individual neurons, and the light blue arrows the apical dendrites of the same neurons. (**B**) Detail of the cerebral cortex of the same brain (image acquired using a ClearScope as described in (A)); the image was deconvolved using the Neuro Deblur™ software (MBF Bioscience)^20^. The arrows indicate a cell process that passes tangentially through this part of the cerebral cortex and can be traced over a distance of 650 µm in this image. The scale bar in (A) represents 250 µm and the scale bar in (B) 100 µm.

**Fig. 11.**
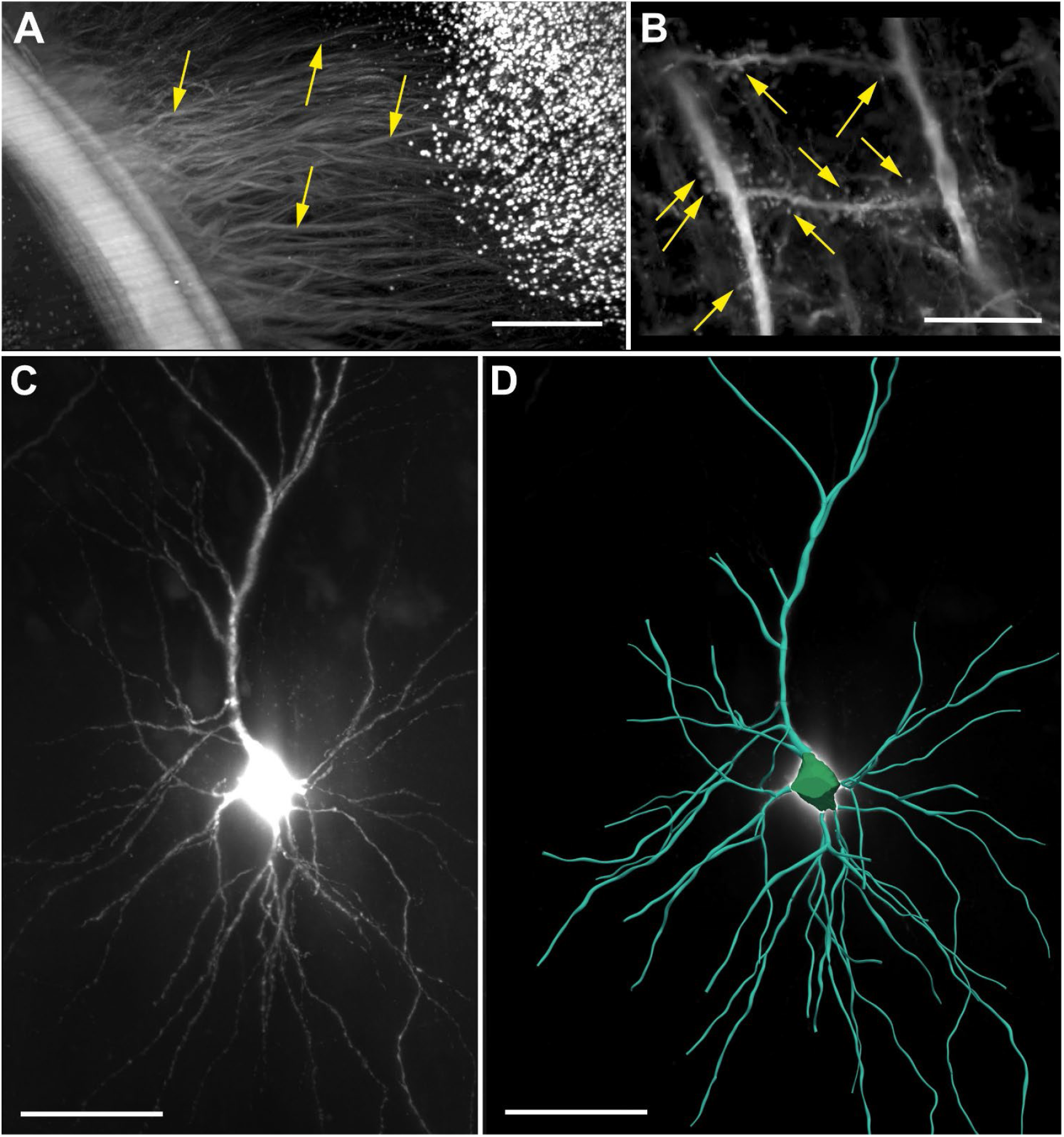
(**A**) The subcortical white matter region indicated by the yellow rectangle in Fig. 9C shown at higher magnification (of the brain from a transgenic mouse expressing eGFP under control of the Thy-1 promoter (hereafter, Thy1-eGFP)), acquired using a ClearScope (developed in this project) with a MBF Bioscience modified UPLFLN4XPH objective (4x/0.13 NA; modified WD = 17 mm; Olympus), 488 nm excitation laser, ZET405/488/561/640mv2 quad band emission filter (Chroma Technologies) and Prime BSI Express camera (Teledyne Photometrics). The arrows point to cell processes that can be individually traced. (**B**) High-magnification maximum intensity projection of a portion of a 3D image of the brain of a transgenic mouse expressing Thy1-eGFP. The brain was cleared using CLARITY^5-7^ before imaging; the 3D image was captured using a ClearScope with a CFI90 20XC Glyc objective (20x/1.0 NA ∞, WD = 8.2 mm; Nikon), 488 nm excitation laser, ZET405/488/561/640mv2 laser quad band emission filter (Chroma Technologies) and Orca-Flash camera in V3 mode (Hamamatsu Photonics). The arrows indicate quantifiable dendritic spines of cortical neurons. (**C**) Portion of a human postmortem entorhinal cortex cleared using iDisco^8^; neurons were filled with biocytin dye after patch-clamping and subsequent staining using streptavidin tagged with Alexa 555. The 3D image was captured using a ClearScope with a CFI90 20XC Glyc objective (Nikon), 561 nm excitation laser, ZET405/488/561/640mv2 laser quad band emission filter (Chroma Technologies) and Orca-Flash camera in V3 mode (Hamamatsu Photonics). (**D**) Neuron from (**C**) digitally reconstructed for quantitative analysis using the automatic 3D neuron reconstruction software, Neurolucida 360^18^. The scale bar in (A) represents 500 µm and the scale bars in (B-D) 100 µm each.

Figure 11B shows quantifiable dendritic spines of cortical neurons in a high-magnification MIP of a portion of a 3D image of the brain from a transgenic mouse expressing Thy1-eGFP; the image was acquired using a CFI90 20XC Glyc objective (20x/1.0 NA ∞, WD = 8.2 mm; Nikon).

The same objective was used to image a portion from a human postmortem entorhinal cortex with neurons filled with biocytin dye after patch-clamping and subsequent staining using streptavidin tagged with Alexa 555 (Fig. 11C). The neuron shown in Figure 11C was digitally reconstructed for quantitative analysis using the automatic 3D neuron reconstruction software, Neurolucida^®^ 360 (MBF Bioscience)^18^.

Figure 12A-C shows MIPs of a portion of a 3D image of an entire, intact mouse brain with vessels labeled using *Lycopersicon Esculentum* (Tomato) Lectin (LEL, TL), DyLight 649 (ThermoFisher Scientific) that was acquired using a XLPLN10XSVMP objective (Olympus); Figure 13A provides a 3D representation of the same portion of the 3D image shown in Figure 12A-C. The vessels shown in Figures 12A-C and 13A were digitally reconstructed for quantitative analysis using the automatic reconstruction software, Neurolucida 360 (MBF Bioscience)^18^ (Fig. 12a-c and Fig. 13B).

**Fig. 12.**
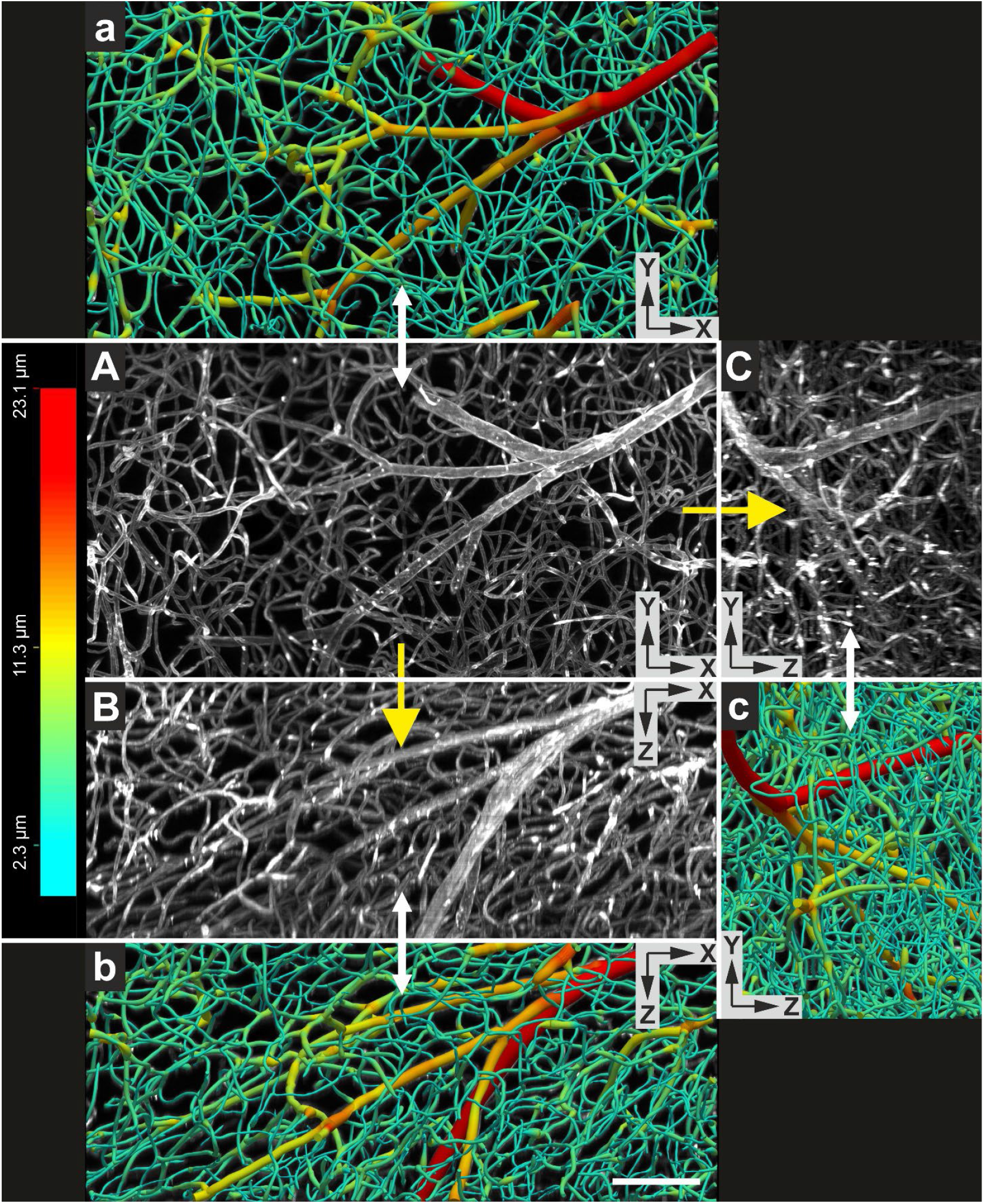
(**A**-**C**) Maximum intensity projections (XY view in (**A**); XZ view in (**B**) and YZ view in (**C**)) of a portion of a 3D image of an entire, intact mouse brain with vessels labeled using *Lycopersicon Esculentum* (Tomato) Lectin (LEL, TL), DyLight 649 (ThermoFisher Scientific) acquired using a ClearScope (developed in this project) with a XLPLN10XSVMP objective (10x/0.6 NA ∞, WD = 8 mm; Olympus), 640 nm excitation laser, ZET405/488/561/640mv2 quad band emission filter (Chroma Technologies) and Prime BSI Express camera (Teledyne Photometrics). The brain was provided by Binaree, Inc. and cleared with Binaree Rapid Tissue Clearing^11^. (**a**-**c**) Vessels in (A-C) digitally reconstructed for quantitative analysis using the automatic 3D reconstruction software, Neurolucida 360^18^. Vessel diameters are color-coded as shown on the left. The scale bar in (b) represents 100 µm.

**Fig. 13.**
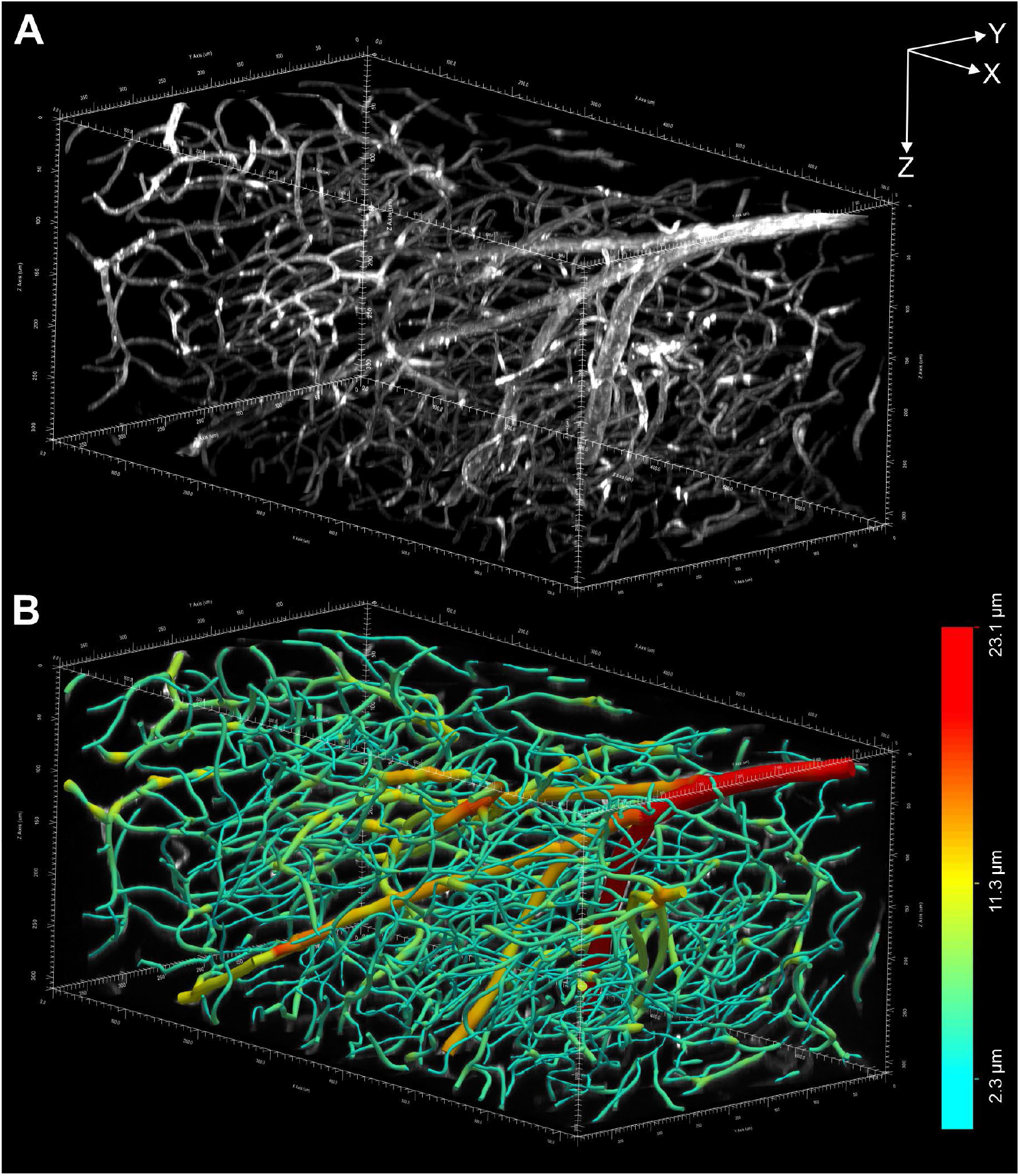
**(A)** 3D representation of the 3D image of an entire, intact mouse brain with vessels labeled using Lycopersicon Esculentum (Tomato) Lectin (LEL, TL), DyLight 649 (ThermoFisher Scientific) acquired using a ClearScope (developed in this project) whose XY, XZ and YZ maximum projections are shown in Fig. 12A-C. **(B)** Vessels in (A) digitally reconstructed for quantitative analysis using the automatic 3D reconstruction software, Neurolucida 36018. The vessel diameters are color-coded as shown on the right. Each of the arrows in (A) represents 100 µm.

### Registration of 3D images acquired using the ClearScope with the Allen Mouse Brain Common Coordinate Framework (CCFv3)

The exceptional image quality of the 3D images acquired from entire, intact mouse brains with the ClearScope (c.f. Fig. 9) enables researchers to view virtual sections through these 3D images and register them with the Allen Mouse Brain Common Coordinate Framework (CCFv3)^21-23^, as shown in Figure 14. This functionality can be used for quantitative analysis as well to perform an initial fast, low-magnification scan, followed by selection of anatomic ROIs to be subsequently scanned at higher magnification, combined with systematic random sampling in LSTM imaging according to the principles of design-based stereology^24,25^. This approach will result in increased imaging speed and further reduced data storage combined with full documentation of image regions that were analyzed.

**Fig. 14.**
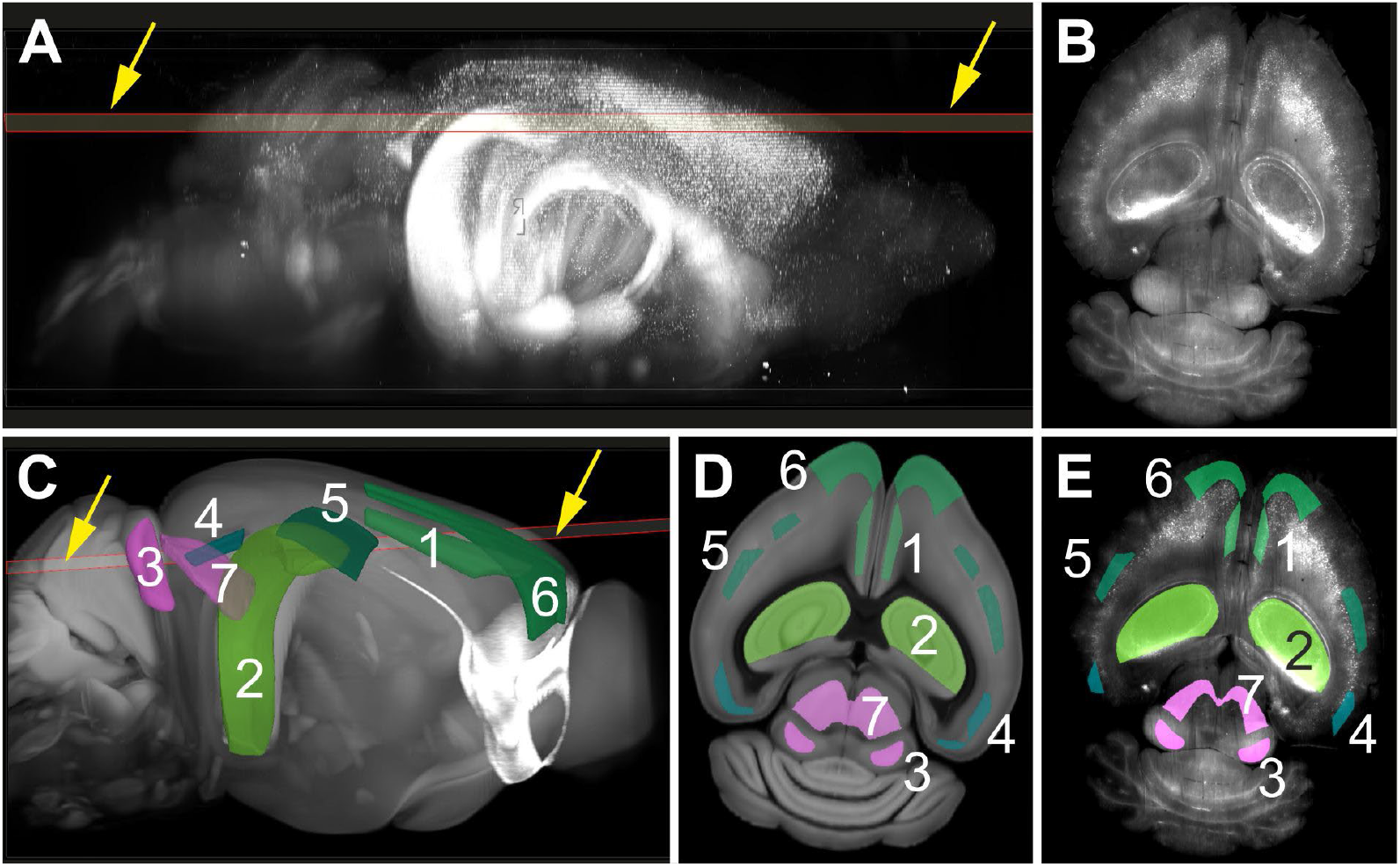
Automatic identification of anatomic regions using the NeuroInfo^®^ software (MBF Bioscience)^22,26^ in a low-magnification scan of brain tissue acquired using a ClearScope (developed in this project) for selective imaging at high magnification and subsequent, automatic or computer-assisted quantitative analysis. (**A**) Selection of a virtual section (arrows) through the 3D image of an entire, intact brain from a transgenic mouse expressing eGFP under control of the Thy-1 promoter, captured using a ClearScope (shown in Fig. 9B) (hereafter: virtual brain section). (**B**) Selected virtual brain section. (**C**) Automatic registration of the volumetric image of the brain, and the selected virtual brain section (arrows) with the Allen Mouse Brain Common Coordinate Framework (CCFv3)^21-23^ using the NeuroInfo software^22,26^. Some anatomic regions featured by the CCFv3 are indicated (1, anterior cingulate area, dorsal part, layer 5; 2, field CA1 hippocampus; 3, inferior colliculus, central nucleus; 4, lateral visual area, layer 5; 5, primary somatosensory area, barrel field, layer 4; 6, secondary motor area, layer 2-3; 7, superior colliculus, motor related, intermediate white layer). (**D**) Virtual section through the CCFv3 with indication of the anatomic regions depicted in (C), representing the best match to the virtual mouse brain section shown in (A,C). (**E**) Automatic delineation of the anatomic regions identified in (C,D) in the virtual mouse brain section selected in (A,C).

## DISCUSSION

There is a growing need for microscopes that are capable of imaging intact organs at high resolution. The initial application for the ClearScope is in the field of neuroscience research. The example photomicrographs shown in Figs 9-11 demonstrate the immense potential of the ClearScope in connectomics research. In the example shown in Fig. 9 the processing time (including fixation and clearing) was approximately 10 days; imaging took 8 hours. In addition to the field of neuroscience researching the central nervous system, there are also applications of the ClearScope in the field of systems biology to better understand innervation patterns in peripheral organs, cancer research to better understand angiogenesis patterns in various organs that are affected by tumors, and analysis of organoids.

While there is no light sheet microscope available with the image quality and unlimited lateral extent of the ClearScope, there are numerous commercial light sheet microscopes on the market. Table 2 (modified from^27^) summarizes and compares the key technical specifications (detection objectives with numerical aperture, axial resolution, refractive index range, maximum depth and maximum specimen lateral size) of commercially available light sheet microscopes, as well as of the recently developed hybrid open-top light-sheet (Hybrid OTLS) microscope ^27^. Figure 15 shows the principles of imaging with the light sheet microscopes summarized in Table 2.

**Table 2.** Key technical specifications of commercially available light sheet microscopes and the recently developed, non-commercial Hybrid OTLS (data taken in part from^27^). Abbreviations: Ref, reference; DetObj, detection objectives; NA, numerical aperture; AxRes, axial resolution; RIR, refractive index range; MD, maximum depth; MSLS, maximum specimen lateral size; *, information not provided by the manufacturer; ** higher magnification is achieved with a motorized 16:1 zoom in the detection light path by means of a Zeiss Axio Zoom.v16 (Carl Zeiss Microscopy, Oberkochen, Germany). Details are in the text.

| System | Ref | DetObj | AxRes<br>($\mu\text{m}$) | RIR | MD (mm) | MSLS (X Y, mm) |
| --- | --- | --- | --- | --- | --- | --- |
| Alpha3<br>(PhaseView, Seattle, WA,<br>USA)<br>(Fig. 15A) | [28] | 2x (NA = 0.14)<br>4x (NA = 0.28)<br>10x (NA = 0.50) | 2.0 – 12.0 | 1.33 – 1.56 | 15 | 15 x 15 (standard)<br>15 x 25 (extended) |
| Blaze<br>(Miltenyi Biotech, Bergisch<br>Gladbach, Germany)<br>(Fig. 15A) | [29] | 1.1x (NA = 0.10)<br>4x (NA = 0.35)<br>12x (NA = 0.53) | 4.0 – 24.4 | 1.33 – 1.56 | 17 (1.1x)<br>16 (4x)<br>10.9 (12x) | 24 x 50 |
| Lightsheet 7<br>(Zeiss, Oberkochen,<br>Germany)<br>(Fig. 15A) | [30] | 5x (NA = 0.16)<br>10x (NA = 0.50)<br>20x (NA = 1.00)<br>40x (NA = 1.00)<br>63x (NA = 1.00) | 2.0 – 14.0 | 1.33 – 1.58 (5x)<br>1.33 (10x)<br>1.33 – 1.53 (20x)<br>1.33 (40x)<br>1.33 (63x) | 10 | 10 x 50 |
| MuVi SPIM CS<br>(Bruker, Billerica, MA, USA)<br>(Fig. 15B) | [31] | 10x (NA = 0.50)<br>20x (NA = 1.00) | 2.0 – 8.0 | 1.33 – 1.51 (10x)<br>1.44 – 1.50 (20x) | 12 | 12 x 19 |
| AxL Cleared Tissue<br>LightSheet (Intelligent<br>Imaging Innovations)<br>(Fig. 15C) | [32] | 1.0 (NA = 0.25)**<br>1.5 (NA = 0.37)**<br>2.3 (NA = 0.57)** | * | 1.33 – 1.56 | 56 (1.0x)<br>30 (1.5x)<br>10 (2.3x) | 100 x 30 |
| SmartSPIM<br>(Lifecanvas, Cambridge, MA,<br>USA)<br>(Fig. 15C) | [33] | 3.6x (NA = 0.20)<br>15x (NA = 0.40) | 1.4 – 4.0 | 1.33 – 1.56 | 12 | 20 x 25 (standard)<br>30 x 55 (extended) |
| ct-dSPIM<br>(ASI, Eugene, OR, USA)<br>(Fig. 15D) | [34] | 16x (NA = 0.40)<br>24x (NA = 0.70) | >1.0 | 1.33 – 1.56 | 5 (16x)<br>2 (24x) | Unconstrained |
| CTLS<br>(Intelligent Imaging<br>Innovations, Denver, CO,<br>USA)<br>(Fig. 15D) | [35] | 1x (NA = 0.20)<br>1.5x (NA = 0.37) | 3 | 1.33 – 1.56 | 25 | 25 x 25 |
| MegaSPIM (Lifecanvas)<br>(Fig. 15D) | [36] | 1.8x*<br>3.6x (NA = 0.20)<br>9x*<br>15x (NA = 0.40)<br>22x* | * | * | * | 200 x 200 |
| Hybrid OTLS (open-top)<br>(Fig. 15F/G) | [27] | 2x (NA = 0.10)<br>24x (NA = 0.70) | 2.9 – 15.6 | 1.33 – 1.56 | 10 | Unconstrained |
| ClearScope<br>(MBF Bioscience)<br>(Fig. 15E) |  | 4x (NA = 0.28)<br>10x (NA = 0.60)<br>16x (NA = 0.40)<br>20x (NA = 1.00)<br>24x (NA = 0.70) | 2.0 – 6.0 | 1.33 – 1.56 | 17 (4x)<br>8 (10x)<br>12 (16x)<br>8 (20x)<br>10 (24x) | Unconstrained |

**Fig. 15.**
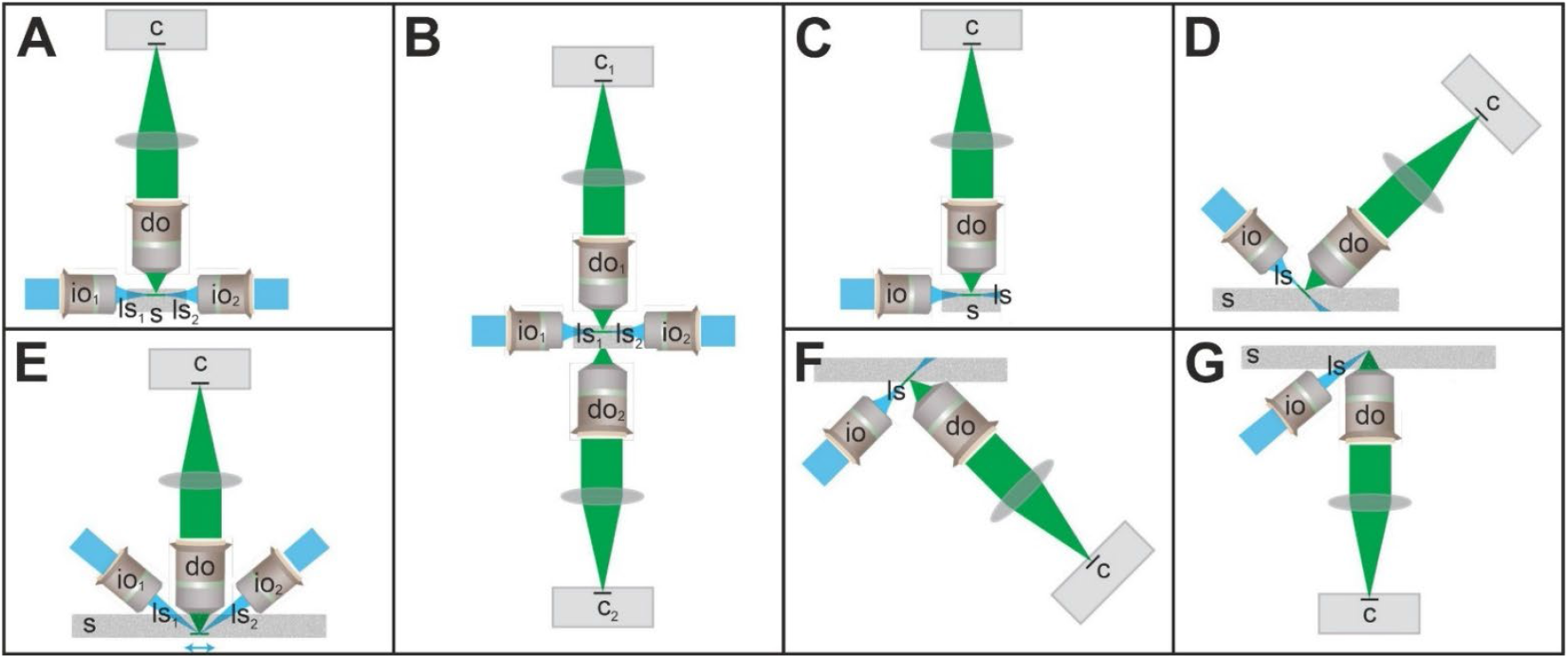
Principles of imaging with the light sheet microscopes summarized in Table 2. Abbreviations: c, camera; do, detection objective; io, illumination objective; ls, light sheet; s, specimen. Details are in the text. The panels were modified from^13^ published under the CC BY 4.0 license (https://creativecommons.org/licenses/by/4.0/).

The classical Selective Plane Illumination Microscopy (SPIM) design principle of imaging with one, two or more light sheets^37,38^ (Fig. 15A) is realized in the Blaze (Miltenyi Biotech), Alpha3 (Phase View) and Lightsheet 7 (Zeiss). However, this design principle results in constrained lateral imaging capabilities (c.f. Table 2). The MuVi SPIM CS (Bruker) uses a modified SPIM design principle with two light sheets and two detection objectives (Fig. 15B), resulting in the same constraints in lateral imaging capabilities.

The SmartSPIM and MegaSpim (both from Lifecanvas) as well as the ct-dSPIM (ASI), CTLS and AxL Cleared Tissue (both from Intelligent Imaging Innovations) use the Axially Swept Light-Sheet Microscopy (ASLM) technology^39^ (Fig. 15C,D). The ASLM technology can be considered a precursor of the LSTM technology and makes use of only one light sheet. This light sheet is axially swept over the field-of-view within a certain focal plane inside the specimen, and emission light is collected using a rolling shutter. The angle between the long axis of the light sheet and the long axis of the detection objective is restricted to 90 degrees, as any angle between the long axis of the light sheet and the long axis of the detection objective less than 90 degrees is protected by Dr. Tomer’s LSTM patent^40^. The long axis of the detection objective can be oriented perpendicular to the surface of the specimen (Fig. 15C) as in the SmartSPIM (Livecanvas) and the AxL Cleared Tissue (Intelligent Imaging Innovations), resulting in constrained lateral imaging capabilities. Alternatively, the long axis of the detection objective can be oriented e.g., in a 45-degree position relative to the surface of the specimen (Fig. 15D) as in the MegaSPIM (Lifecanvas) and the ct-dSPIM (ASI), resulting in constraints in the maximum depth that can be imaged.

Unconstrained lateral imaging is only possible with the ClearScope (Fig. 15E), the ct-dSPIM (ASI) and the Hybrid OTLS^27^ (Fig. 15F,G).

Compared to the ct-dSPIM (ASI) the ClearScope provides low-magnification (4x and 5x) objectives in addition to high-magnification objectives (10x, 20x, 24x, 27x), and a maximum penetration depth that is more than twice as high when using a 16x detection objective (12 mm vs 5 mm) and five times higher when using a 24x detection objective (10 mm vs 2 mm).

The MegaSPIM (Lifecanvas) offers a maximum specimen lateral size of 200 mm x 200 mm, which is suitable for imaging large, cleared thick tissue slabs from human and non-human primate postmortem brains. However, the long axis of the detection objective is oriented in a 45-degree position relative to the surface of the specimen and, thus, compared to the ClearScope, the maximum depth that can be imaged with the MegaSPIM is WD x sin (45°), with WD = working distance of the detection objective. Furthermore, an example 3D image^36^ provided by the manufacturer of the MegaSPIM (Lifecanvas) (showing phospho-tau and alpha-synuclein in a 1 mm-thick coronal section of a human postmortem brain, imaged with a MegaSPIM and a 1.8x detection objective) demonstrated a vertical and horizontal banding pattern that is not ideal for quantitative analysis, in contrast to the seamless 3D images obtained with the ClearScope (c.f. Fig. 9).

The Hybrid OTLS^27^ is the only open-top light sheet microscope in this list. For imaging at low magnification, the Hybrid OTLS^27^ uses a conventional (orthogonal) architecture (Fig. 15F), and for imaging at high magnification a non-orthogonal architecture (Fig 15G) (note that the later prevents commercialization of the Hybrid OTLS^27^ as any angle between the long axis of the light sheet and the long axis of the detection objective less than 90 degrees is protected by Dr. Tomer’s LSTM patent^40^).

The developers of the Hybrid OTLS specified the following five key requirements for next-generation light sheet microscopes:^27^ (i) user-friendly mounting of multiple specimens with standard holders, (ii) compatibility with all current clearing protocols, (iii) no fundamental limits on lateral specimen size, (iv) a large imaging depth to accommodate intact mouse organs and thick tissue slabs, and (v) broad ‘multi-scale’ imaging capabilities for time- and data-efficient workflows. All these requirements are achieved in the design of the ClearScope.

Of note, compared to the ClearScope the Hybrid OTLS^27^ has a number of disadvantages: (i) a much more complex illumination path; (ii) support of only 4 laser wavelengths compared to 7 laser wavelengths supported by the ClearScope; (iii) only two detection objectives (2x and 24x), with lower axial resolution than the detection objectives used in the ClearScope (c.f. Table 2); (iv) no possibility to change the detection objectives according to the specific needs of researchers; and (v) the low magnification (2x) detection objective oriented in a 45 degree position relative to the surface of the specimen, which reduces the maximum depth that can be imaged with this objective to 10 mm (compared to 12 mm thickness that can be imaged with the low magnification (4x) objective of the ClearScope).

It should be mentioned that Glaser et al.^27^ stated in their publication describing the Hybrid OTLS that synchronizing the two light sheets in LSTM (and, thus, in the ClearScope) to a narrow confocal slit (rolling shutter) would result in “inefficient light collection” and “fundamental losses introduced by the LSTM scanning strategy” (Supplementary Note 1 in^27^). Unfortunately, this statement fully disregards all the proven advantages of implementing a line-scan imaging strategy in light sheet microscopy that are realized in both the ASLM technology^39^ and the LSTM technology^13^.

In addition to these commercially available light sheet microscopes, the OpenSPIM should be mentioned; it is an open source SPIM imaging option.^41,42^ According to^43^ one can construct a working SPIM system from commercially available components and 3D-printed or custom-machined parts. The system is capable of imaging smaller specimens and provides scanning capabilities using low precision positioning devices. Speed is very slow and requires a high degree of mechanical aptitude from the user for assembly and alignment. Neither the original OpenSPIM^41^ nor the more recent X-OpenSPIM^42^ offer unconstrained lateral imaging.

Early adopters in research labs have acquired the ClearScope system and published cutting-edge biomedical research,^44-46^ with microscopy capabilities that were not previously possible. For example, Datta et al.^44^ repeatedly administered the anesthetic, pain medication and fast-acting antidepressant ketamine to mice and performed whole-brain imaging of dopamine neurons (sample size after clearing was ∼ 15 x 20 mm, and 9 mm deep) using a ClearScope with 4x and 20x objectives. The authors found a dosage-dependent decrease in the number of dopamine neurons in the behavior state-related midbrain regions, a dosage-dependent increase in the number of dopamine neurons within the hypothalamus, and divergently altered innervations of prefrontal cortex, striatum, and sensory areas.^44^ As per Table 2 this research would not have been possible with any other commercially available light sheet microscope without cutting the mouse brains into pieces, which would negatively impact reconstruction of the connectivity pattern within the brain. Of note, hypothetically, Datta et al.^44^ could have alternatively used the Hybrid OTLS microscope^27^ in their research. However, the Hybrid OTLS is not available commercially.

The development of the ClearScope addresses two fundamental requests by experts in the field:^47^ (i) to focus on practicality and applicability to biological and biomedical research questions in further developing light sheet microscopy, rather than focusing on optimizing technical specifications of light sheet microscopes to their maximum extent (such as the numerical aperture that can be covered in light sheet microscopy^48,49^) that would likely also diminish the practicability of the instrument; and (ii) to develop smart imaging schemes that explore specimens at low magnification and autonomously switch to higher magnification imaging only in areas of interest.^47^ The resulting dataset sizes will be dramatically reduced in comparison to current methods of scanning the entire specimen at the magnification required for analysis.

### Limitations

One limitation of this study is that the LSTM technology (and, thus, the ClearScope) requires more light intensity than traditional light sheet microscopes, so it requires more powerful lasers. On the other hand, this is justified considering the exceptional image quality of the ClearScope. The same applies to another limitation, namely the fact that the ClearScope illumination optics are optimized for imaging using high magnification objectives (up to 25x) and that the trade-off is not being able to image using very low magnification objectives (e.g. 2x).

Another limitation is that the ClearScope software does not have an Application Programming Interface (API). In this regard it was stated in a recent review^47^ on light sheet microscopy that it is considered essential in the neuroscience community that manufacturers of commercial microscopy systems offer interfaces for microscope control, imaging and image analysis using open-source software (e.g. Micro-Manager^50^, Pycro-Manager^51^, AutoScanJ^52^, ImSwitch^53^, MicroMator^54^ and other, Python-based control software^55,56^. It cannot currently be assessed whether only a few scientists or larger parts of the neuroscience community would like to have APIs for control of commercial microscopy systems. If this request is made to us more frequently, we will generate a Python-based API for the ClearScope using Pybind11^57^, a lightweight, header-only library that exposes existing C++ types in Python and vice versa. Pybind11^57^ enables seamless interoperability between C++ and Python, leveraging the performance and efficiency of C++ while providing the flexibility and ease-of-use of Python. This will be achieved by creating Python bindings for the C++ modules of the ClearScope software, allowing them to be imported and used directly within Python scripts.

## CONCLUSIONS

We are convinced that the development of the ClearScope will open significant new avenues of research by enabling researchers to investigate scientific questions and mechanisms of neuropsychiatric, neurological and other biological disorders that have not been previously considered due to the current limitations in performing high resolution light microscopic imaging of intact, cleared specimens. As such this project will enable progress in developing novel prevention and treatment strategies to combat various neuropsychiatric and neurological disorders, as well as cancer.

## METHODS

The ClearScope was developed as a turn-key product in a compact single-box design, designed to be shipped and installed easily in any lab. The ClearScope is, where possible, self-aligning to enable “lights out” operation.

The ClearScope hardware consists of several sub-assemblies: two illumination arms, the detection arm, the focus column, system base, motorized stage and system enclosure. This modular assembly of the ClearScope hardware facilitates manufacturability. Some of the sub-assemblies were outsourced to contractors with specific skills and expertise. For example, the illumination arm alignment was outsourced to an optical assembly company. The hardware is designed so that shipping brackets can be used to immobilize and brace the system for shipping.

The software for the ClearScope is a fully-integrated desktop application for Windows 11 (64-bit) written in C++. It uses MBF Bioscience’s Core Software Libraries and the robust object-oriented design methodology for image acquisition and processing already used in all MBF Bioscience’s commercial software products^58^. The software was developed using the Microsoft Visual Studio Professional integrated development environment which provides tools for interactive development, debugging and code performance analysis. The software was profiled and optimized for memory, processor and GPU usage where necessary. Multiple processing threads were used to provide a high-performance application. A significant amount of effort was spent creating a compelling user experience, including creating fully-featured windows, menus and toolboxes that integrate all software components developed. Usability studies were conducted, and a complete product documentation and support tools were created. A fully documented user’s guide was created.

The product validation and usability studies using the different microscope hardware and software components of the ClearScope (as demonstrated and described in Figs 9-14) were performed in close collaboration with Dr. Tomer. To this end, Dr. Tomer provided superfluous material from ongoing research projects in his lab. Furthermore, the images shown in Figs 9, 10, 11A and 12-14 were acquired from mouse brains that were provided by Binaree, Inc. (Daegu, Republic of Korea). Hence, no specific experiments were performed in the framework of developing the ClearScope and no specific IACUC approval was necessary.

The point spread function estimates were performed as described in the *Results* section. Imaging and analysis were performed at the MBF Bioscience office.

## DATA AVAILABILITY

The datasets used and analyzed during the current study are available from the corresponding author on reasonable request. The ClearScope is commercially available.

## AUTHOR CONTRIBUTIONS

Conceptualization: M.F., P.L., N.O’C., S.A., P.A., R.T., J.G.; Methodology and Data curation: M.F., D.D., N. O’C., B.H., J.B.1, N.R., A.W., P.A., R.T.; Visualization: M.F., N.O’C., A.W., J.G.; Resources: M.F., P.L., D.D, N.O’C., B.H., J.B.1, N.R., A.W., S.A., P.A., J.B.2, T.B., C.G., R.T., J.G.; Writing – original draft: J.G., Writing – review & editing: M.F., P.L., D.D, N.O’C., B.H., J.B.1, N.R., A.W., S.A., P.A., J.B.2, T.B., C.G., R.T.; Funding acquisition: J.G.

## NIH GRANT SUPPORT

Research reported here was supported by the National Institute of Mental Health of the National Institutes of Health under award number R44MH116827 to MBF Bioscience and R.T. (Glaser PI). The content is solely the responsibility of the authors and does not necessarily represent the official views of the National Institutes of Health.

NYU CTSI Pilot NIH/NCATS UL1TR001445 (Cronstein PI) (JB/OD/TB)

## OTHER FUNDING

Parekh Center for Interdisciplinary Neurology (PCIN) Pilot Research Grant (JB/OD)

NYU FACES (TB/JB/OD)

American Epilepsy Society Junior Investigator Award (JB)

## COMPETING INTERESTS

M.F., P.L., D.D., N.O’C., B.H., J.B., N.R., A.W., S.A. and P.A. are employees of MBF Bioscience (Williston, VT, USA). J.G. is founder, president and major shareholder of MBF Bioscience. R.T. served as paid consultant for MBF Bioscience during the first (feasibility) phase of generating the ClearScope.

The LSTM technology is protected by US Patent US11506877B2 (Inventor: Dr. Raju Tomer; Assignee: Columbia University, New York, NY, USA) and related, international patent applications (EP3538941A4, WO2018089839A1 and JP2020502558A). MBF Bioscience has negotiated an exclusive license from Columbia University for the LSTM technology, and a non-exclusive license from the Allen Institute (Seattle, WA, USA) for using the Allen Mouse Brain Common Coordinate Framework (CCFv3)^21,23^.

The name ‘ClearScope’ is protected through copyright by MBF Bioscience.

O.D, J.B., T.B and C.G. declare no competing interests.

## Acknowledgments

Dr. Charles Gerfen provided invaluable neuroanatomical expertise in assessing the accuracy of the registrations of experimental mouse brains to the Allen Mouse Brain Atlas.

